# Identification of Amino Acid Residues Responsible for Differential Replication and Pathogenicity of Avian Influenza Virus H5N1 Isolated from Human and Cattle in Texas, US

**DOI:** 10.1101/2025.03.01.640810

**Authors:** Mahmoud Bayoumi, Ramya S. Barre, Ruby A. Escobedo, Vinay Shivanna, Nathaniel Jackson, Chengjin Ye, Adolfo García-Sastre, Ahmed Mostafa, Luis Martinez-Sobrido

## Abstract

Highly pathogenic avian influenza viruses (HPAIV) pose a serious public health concern. In March 2024, a first-time outbreak of HPAIV H5N1 in dairy cattle herds was reported in the United States (US). Since then, the virus has continued to spread in cattle herds and spilt over into humans. We recently showed that the first human isolate reported in the US in Texas (HPhTX) from a dairy worker in an affected cattle farm has enhanced replication kinetics and pathogenicity in mice compared to a closely related bovine isolate (HPbTX). However, the molecular determinants of differential pathogenicity have not yet been identified. Herein, we show that HPhTX has enhanced polymerase activity, compared with HPbTX, in human cells and that the polymerase basic 2 (PB2) protein is the main factor responsible for this difference. Through single and combined site-directed mutagenesis and swapping the three amino acids different between HPhTX and HPbTX, we found that PB2 mutation E627K is the major contributor to the enhanced polymerase activity of HPhTX. E362G substitution in HPhTX PB2 affected the polymerase, although to a lesser extent than E627K. Moreover, M631L mutation in HPhTX PB2 enhanced polymerase activity. Rescue of a loss-of-function recombinant HPhTX (rHPhTX) containing mutations at residues 627 and 362, alone or in combination, revealed a contribution of PB2 E362G and K627E in morbidity, mortality, and viral replication as compared to rHPhTX wild-type (WT), and significantly reduced viral pathogenicity to levels comparable to rHPbTX WT. These findings indicate that HPAIV H5N1 of cattle origin isolated from the first human case has post-transmission amino acid changes that increase viral replication in human cells and pathogenicity in mice.

## Introduction

Influenza A viruses (IAVs) are enveloped, segmented, single-stranded, negative-sense RNA viruses belonging to the family *Orthomyxoviridae.* Influenza viruses are categorized into influenza A (IAV), B (IBV), C (ICV), and D (IDV), with various host ranges for each type ^1,2^. The natural host reservoirs of most IAVs is wild waterfowl. However, IAVs have the capability to spillover and adapt to infect a broad host range of domestic birds and multiple terrestrial and sea mammals, including humans ^3,4^. The IAV genome is composed of eight RNA segments encoding several essential proteins and various accessory proteins, which differ between strains. IAVs encode three polymerase subunits (PB2, PB1, PA) and the viral nucleoprotein (NP), which encapsidate the viral RNA to form the viral ribonucleoprotein (vRNP) complexes involved in viral genome replication and gene transcription ^1,5,6^.

Since the highly pathogenic avian influenza virus (HPAIV) A/goose/Guangdong/1/1996 (GsGD) H5N1 was first reported in China, the GsGD lineage of HPAIV H5N1 has spread around the world through migratory wild birds, infecting a variety of species and endangering the public health of both humans and animals ^7^. The H5-type GsGD is categorized into clades and subclades according to the H5 gene. The most predominant clade is 2.3.4.4, which first appeared in China in late 2013. Then, the virus spread further, resulting in other outbreak episodes among wild birds and poultry in North America, Europe, and Asia ^8^. Meanwhile, the virus experienced many reassortments with other AIVs, leading to the formation of different subclades (2.3.4.4a–2.3.4.4h) and genotypes of H5Nx subtypes ^9,10^. The most common reassortant subclade in Asia, Europe, and Africa was the HPAIV H5N1 clade 2.3.4.4b in 2021, that exists in different genotypes according to the origin of their six additional genes encoding internal virus proteins ^11^. Several outbreaks among chicken populations occurred in December 2021 as a result of the HPAIV H5N1 spreading from Europe to North America via migrating birds over the Atlantic Ocean ^12^. Shortly after, the HPAIV H5N1 clade 2.3.4.4b arrived in Central and South America, a large number of mammals, including humans and different terrestrial and sea mammals, were also affected ^8,13,14^.

During the first quarter of 2024, an unprecedented outbreak of HPAIV H5N1 clade 2.3.4.4b genotype B3.13 impacting dairy cattle emerged in Texas, New Mexico, and Kansas, rapidly spreading to other states in the US ^15^. So far, the virus has affected nearly 900 herds across approximately 16 states ^15,16^. This outbreak is believed to have originated from a single spillover event from wild birds ^15,16^, followed by sustained transmission between herds, with two more recent introductions from genotype D1.1 taking place in 2025. Contaminated milking equipment and the movement of asymptomatic infected animals and/or contaminated equipment between farms and states, facilitated the spread of the virus ^16,17^.

Bovine infection, thus far, with HPAIV H5N1 typically results in mild mastitis and respiratory issues, with low mortality rates ^15,18^. Beyond cattle, the virus has also affected several other species through direct or indirect contact, including chickens, peridomestic birds, turkeys, alpacas, cats ^16,19^, and humans ^20,21^. Interestingly, ingestion of infected milk has proven fatal in cats ^16,19^. However, human infections with the bovine-origin HPAIV H5N1 have generally been limited to conjunctivitis and/or mild respiratory illness ^20,21^, and the virus has not being able to transmit from human to human.

Recently, we identified that the first human isolate reported in Texas in April 2024 (A/Texas/37/2024 H5N1; hereby named HPhTX) in a dairy worker in an affected cattle farm has enhanced replication kinetics and pathogenicity in C57BL/6 mice compared to the closely related cattle isolate in Texas in April 2024 (A/bovine/Texas/24-029328-02/2024 H5N1; hereby named HPbTX) ^22^. However, the molecular determinants of different replication and pathogenicity between HPhTX and HPbTX have not yet been identified. Herein, we found that HPhTX has enhanced polymerase activity in human cells, compared with HPbTX, and that the polymerase basic 2 (PB2) protein is the main factor responsible for this enhanced polymerase activity. Among the three amino acid differences between HPhTX and HPbTX, we found that PB2 amino acid E627K substitution is the primary factor responsible for differences in polymerase activity. Moreover, PB2 amino acid G362E substitution, but not L631M, also plays a role in the differences in polymerase activity between the human and bovine isolates. Notably, we observed synergistic activity of E627K and G362E in enhancing polymerase activity, replication, and pathogenicity in C57BL/6J mice.

## Materials and Methods

### Biosafety

*In vitro* infections and mice experiments with HPAIV H5N1 were performed at biosafety level (BSL) 3 and animal BSL3 (ABSL3) facilities, respectively, at the Texas Biomedical Research Institute (Texas Biomed). Experiments were reviewed and approved by the Texas Biomed Institutional Biosafety (IBC) and Animal Care and Use (IACUC) committees.

### Cells

Madin-Darby canine kidney (MDCK, ATCC CCL-34), human embryonic kidney (HEK293T, ATCC CRL-3216), Madin-Darby bovine kidney (MDBK, ATT CCL-22), and human lung adenocarcinoma epithelial (A549, ATCC CCL-185) cells were maintained in Dulbecco’s modified Eagle medium (DMEM) (Invitrogen, US) supplemented with 10% fetal bovine serum (FBS) and 1% PSG (i.e., penicillin, 100 U/ml; streptomycin 100 μg/ml; l-glutamine, 2 mM) at 37°C in 5% CO_2_ incubators. MDBK cells were kindly provided by Professor Daniel Perez at the University of Georgia, Athens, GA, US.

### Viruses

Recombinant human A/Texas/37/2024 H5N1 (rHPhTX) and bovine A/bovine/Texas/24-029328-02/2024 (rHPbTX) wild-type (WT) viruses were generated as we described earlier ^22^. Briefly, the viral segments of HPhTX (GenBank accession # PP577940-47) were synthesized based on published sequences and subcloned in the pHW2000 plasmid. The segments of HPbTX (GenBank accession # PP599470.77), which exhibit variations compared to HPhTX, underwent site-directed mutagenesis (SDM) to generate the corresponding rHPbTX ^22^.

### Rescue of rHPhTX PB2 mutants

To determine the impact of PB2 mutations on viral replication and pathogenicity, we generated various PB2 mutants in the pHW2000 plasmid by SDM with primers listed in **Table 1** using the QuikChange Mutagenesis kit (Agilent, US) according to the manufacturer’s instructions. The rHPhTX PB2 mutant viruses were generated as previously described with the pHW2000 plasmids containing the indicated mutations ^22^. Briefly, pHW2000 plasmids (1 μg each) encoding the viral PB2 mutants (PB2-362G, PB2-627E, and PB2-362G/627E), PB1, PA, HA, NP, NA, M and NS segments of HPhTX in serum-free Opti-MEM media were co-transfected into a co-culture of HEK293T and MDCK cell lines using the Lipofectamine™ 3000 Transfection Reagent according to the manufacturer’s recommendations (ThermoFisher Scientific, US). The transfection mixtures were then replaced with 2 ml of Opti-MEM media containing 1% PSG and 0.2% bovine serum albumin (BSA) and incubated for 48 h at 37°C in a 5% CO_2_ cell culture incubator. At 72 h post-transfection (hpt), cell culture supernatants were collected and cleared by centrifugation at 2,500 rpm for 5 min at 4°C. For propagating the rescued viruses, 500 μl of the collected cell culture supernatants were used to infect fresh monolayers of MDCK cells in T-75 flasks with DMEM media containing 1% PSG and 0.2% BSA. The recombinant WT and mutant viruses were aliquoted and stored at −80°C until their use.

**Table 1.**
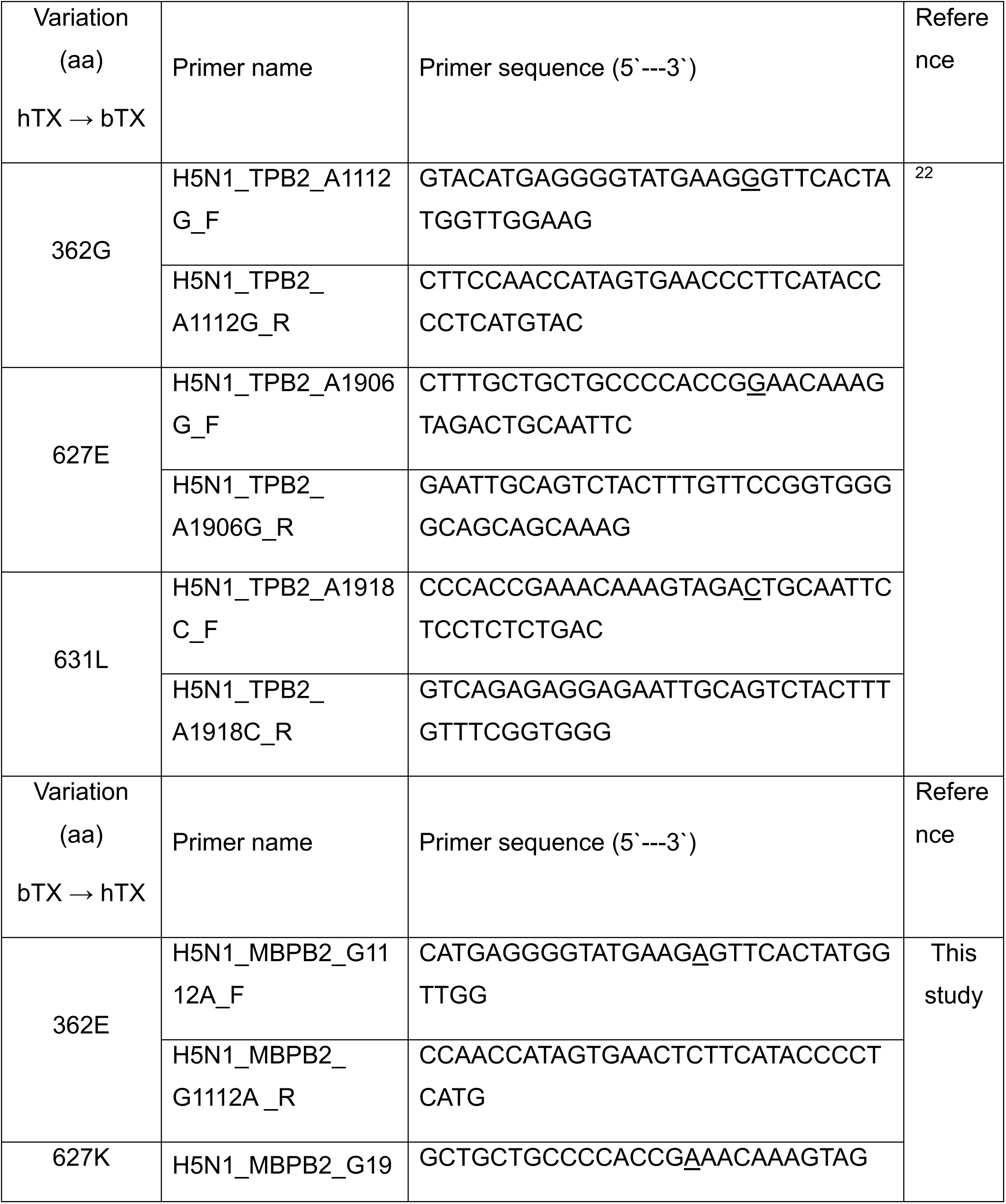
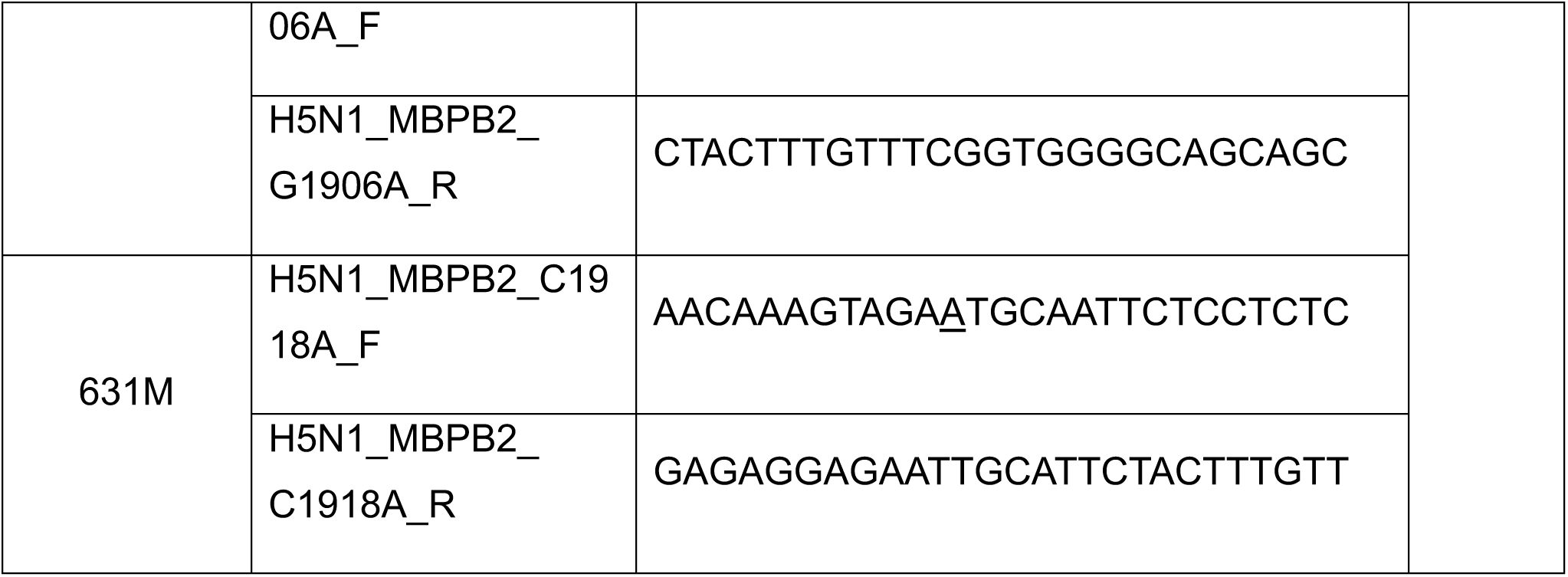
Primers used to generate rHPbTX and rHPhTX PB2 mutants.

### Antibodies

Mouse monoclonal antibodies to detect HPAIV H5N1 polymerase expression were obtained from BEI resources: Clone 170-3C12 was used to detect PB2, Clone F5-46 was used to detect PB1, and Clone 1F6 for detection of PA. A mouse monoclonal antibody (HT103) was used to detect the viral NP (Kerafast, US) in Western blotting assays. A rabbit polyclonal antibody against the viral NP was utilized for murine lung and brain tissue in immunohistochemical staining (ThermoFisher Scientific, US). Monoclonal anti-β actin clone AC-15 (Sigma, US) was used to detect β actin as loading control for western blotting assays.

### RNA extraction, RT-PCR, and sequencing

Viral RNA extraction from rescued WT or mutant viruses was performed according to the TRIzol-LS reagent manufacturer’s instructions (Invitrogen, US). SuperScript^TM^ II Reverse Transcriptase (ThermoFisher Scientific, US) was used to prepare the cDNA synthesis step. PCR was conducted with PfuTurbo DNA Polymerase (Agilent, US) using the following primers: PB2 forward, 5’-TATTGGTCTCAGGGAGCGAAAGCAGGTC-3’, PB2 reverse, 5’-ATATGGTCTCGTATTAGTAGAAACAAGGTCGTTT-3’ ^23^. The amplification protocol was 95°C for 5 min followed by 40 cycles at 95°C for 10 sec, 58°C for 30 sec, and 72°C for 150 sec. Purified PCR products were subjected to next-generation sequencing (NGS) to verify the desired mutations.

### Viral infections and replication kinetics

Monolayers of A549, MDBK, and MDCK cell lines were cultured in 6-well plates (10^6^ cells per well, triplicates) and infected with the indicated viruses at a multiplicity of infection (MOI) of 0.0001 and kept at 37°C in a 5% CO_2_ incubator to allow viral adsorption for 1 h with gentle tilting every 15 min. Following viral adsorption, the infected cell monolayers were washed twice with PBS to remove residual non-adsorbed viral particles, and plates were then incubated with 1 ml DMEM media containing 1% PSG and 0.2% BSA at 37°C in a 5% CO_2_ incubator. Cell culture supernatants were collected at 12, 24, 48, and 72 h post-infection (hpi). Viral titers were determined using standard plaque assay.

### Quantification of HPAIV H5N1 by plaque assay

Cell culture supernatants collected from infected cells were serially 10-fold diluted and used to infect MDCK cells in 6-well plates (10^6^ cells per well, triplicates). After 1 h of adsorption with gentle tilting every 15 min, the plates were washed twice with PBS. DMEM containing 1% PSG, 0.2% BSA, and warm 1% agarose solution was overlaid immediately after viral infection and before solidification. Plates were incubated at 37°C in a CO_2_ incubator. After 48 h, cells were fixed overnight with a 10% neutral buffered formalin solution to inactivate the virus. Then, the agarose plug was removed and 2 ml of 0.2% crystal violet solution was added to each well and incubated for 20 min at room temperature. Then, the crystal violet solution was removed, and the wells were washed with water to rinse the excess stain solution. The plates were scanned and photographed using a ChemiDoc MP Imaging System.

### Minigenome (MG) assays

To evaluate the impact of WT and mutant viral polymerase proteins on viral genome replication and gene transcription, MG assays were performed as previously described ^24–27^. Briefly, HEK293T cells (12-well plate format, 5×10^5^ cells/well, triplicates) were transiently co-transfected, using Lipofectamine™ 3000 Transfection Reagent, with 1 µg of each indicated pHW2000 plasmids encoding the viral polymerase proteins (PB2, PB1, PA), and NP, together with 1 µg of a MG plasmid encoding an influenza viral RNA (vRNA)-like segment expressing ZsGreen (ZsG) fused to nanoluciferase (Nluc) flanked by the NP segment non-coding regions (NCR). The expression of ZsG-Nluc is driven by a human RNA polymerase I promoter (hPolI) and a mouse RNA polymerase I terminator (mTER). A Cypridina luciferase (Cluc)-expressing plasmid (1 µg) under the control of the chicken β-actin promoter in a pCAGGS plasmid was included to normalize transfection efficiencies ^24–26^. HEK293T transfected cells without pHW2000 PB1 plasmid served as a negative control. At 6 hpt, media was substituted by fresh DMEM media containing 1% PSG and 10% FBS. At 30 hpt, Nluc and Cluc expression levels were determined using a Nano-Glo® luciferase assay system (Promega, US) and a Cypridina luciferase glow assay kit (Thermo Scientific, US), respectively. Promega™ GloMax® luminometer was used to measure luminescence activity. Polymerase activity was calculated by dividing Nluc activity by the Cluc activity and expressed as fold induction compared to the negative control sample. At 30 hpt, representative live cell images showing ZsG expression from cells subjected to various MG assays, as indicated above, were imaged through fluorescence microscopy (EVOS).

### Western blots

Cell monolayers were washed with ice-cold PBS and treated with ice-cold NP40 lysis buffer (Thermo, US; completed with protease inhibitors cocktail). Cells were lysed on ice for 30 min, scrapped off and transferred to a microfuge tube. Cells were centrifuged at 15,000 rpm for 15 min at 4°C. The supernatant of cell lysates was mixed with 2x working LDS sample buffer completed with 20% β-mercaptoethanol and boiled at 98°C for 5 min before loading into SDS-PAGE gels. After running, gels were transferred into nitrocellulose membranes, and membranes were incubated in a blocking buffer (5% non-fat dry milk powder in PBS with 0.05% tween 20 (PBST) (Sigma, US). Blocked membranes were probed with the designated primary antibodies overnight at 4°C. Then, the membranes were washed 3x for 5 min with 5 ml PBST. After washing, the membranes were incubated with secondary antibodies for two hours at room temperature. After 3x washes for 5 min with 5 ml PBST, chemiluminescence-generated protein bands were visualized using a SuperSignal (ThermoFisher, US) ECL substrate kit following manufacturer’s recommendations.

### Mice experiments

C57BL/6J 6-week-old female mice (n=13/group) were anaesthetized intraperitoneally (i.p.) using a cocktail of Xylazine (20 mg/ml) and Ketamine (100 mg/ml). Anaesthetized mice were infected intranasally (i.n.) with the indicated viral doses in 50 μl of DMEM containing 1% PSG and 0.2% BSA. At 2- and 4-days post-infection (DPI), four mice from each group were euthanized to collect lungs, nasal turbinate, and brains. Half of the organs were fixed in a 10% neutral buffered formalin solution for histopathology and immunohistochemistry analyses, and the other half was homogenized in 1 ml of PBS using a Precellys tissue homogenizer (Bertin Instruments, France) for viral titration. Tissue homogenates were centrifuged at 10,000 rpm for 10 min, and the supernatants were used to determine viral titers using standard plaque assay in MDCK cells. The remaining 5 animals in each experimental group were monitored for 14 days for disease progression, body weight changes, and survival. Mice that experienced a weight loss exceeding 25% of their original weight were humanely euthanized.

### Histopathology and immunohistochemistry

At necropsy, mouse lung and brain tissues were collected and fixed in 10% neutral buffered formalin, then paraffin-embedded, sectioned, and stained with hematoxylin and eosin (H&E) as previously described ^22^. Briefly, formalin-fixed tissues were processed in a Tissue-Tek VIP tissue processor that involves dehydration through graded alcohols, clearing with a 50:50 absolute alcohol/xylene mixture and xylene, and paraffin wax embedding using ParaPro™ XLT (StatLab, US). Embedded Paraffin blocks were sectioned at 4 µm using a Microm® HM325 rotary microtome, mounted on glass slides using a 46-48°C water bath, and stained using a Varistain Gemini automated stainer. Deparaffinization steps include xylene, absolute alcohol, and 95% alcohol washes. Hematoxylin (StatLab, US) staining was followed by High-Def solution (StatLab, US) for excess stain removal and Reserve Bluing Reagent (StatLab, US). Eosin (StatLab, US) staining was followed by dehydration through alcohol, an alcohol/xylene mixture, and xylene before cover slipping. A board-certified veterinary pathologist examined the stained sections using light microscopy. All stained slides were scanned at 20x magnification in the Axio Scan Z1 (Zeiss, Germany), and the images were analyzed using HALO software (Indica Labs, US) to calculate percentage pathology using pathologist-developed AI classifier modules. For immunohistochemistry, 4 µm sections were mounted on positively charged slides, air-dried overnight, and processed on a Ventana Discovery Ultra IHC/ISH automatic stainer. Deparaffinization was done using Discovery Wash (Roche, US) and cell conditioning with Discovery CC1 (Roche, US) at 95°C for 64 min. Endogenous peroxidase blocking was performed using Discovery Inhibitor (Roche, US) for 8 min. Sections were incubated with Influenza A NP rabbit polyclonal antibody (ThermoFisher Scientific, US) at 1:1500 dilution for 1 h at room temperature, detected using anti-rabbit HQ (Roche, US) and Anti-HQ HRP (Roche, US) for 8 min each at 36°C, and visualized using ChromoMAP DAB (Roche, US). Hematoxylin (Roche, US) and Bluing Reagent (Roche, US) were used for counterstaining.

### Molecular modelling and prediction

Three-dimensional structure prediction was performed using the PHYRE2 web server ^28^ and I-Tasser ^29^ to predict each representative PB2 with the intensive mode. The PyMOL 3.1.3 (The PyMOL Molecular Graphics System, Version 3.1.3 Schrodinger, LLC) was used to annotate and visualize the known crystalized PDB: 6QNW; H5N1 and PDB: 5wl0; H3N2 as well as predicted bovine PB2 H5N1 structures.

### Quantification and statistical analysis

When multiple comparisons were required for a single factor, the experimental means were compared using a one-way analysis of variance (one-way ANOVA) with Dunnett’s multiple comparison test. Multiple comparisons among different time points and experimental means were compared using a two-way analysis of variance (two-way ANOVA) with Greenhouse-Geisser correction, followed by Tukey’s multiple comparisons analysis. p values were calculated with GraphPad Prism 8 (GraphPad Software, US; www.graphpad.com). The data represents the average of three biological replicates with the standard deviation (SD). ns: non-significant; p>0.05, *p < 0.01, **p < 0.001, ***p < 0.001, ****p < 0.0001.

## Results

### HPhTX has enhanced polymerase activity in human cells compared with HPbTX

We and others have identified that the first human isolate that was reported in Texas, US (HPhTX) from a dairy cattle worker in an affected cattle farm has enhanced replication kinetics and pathogenicity in mice compared to closely related cattle isolates in Texas, US, in April 2024 (HPbTX) ^22,27^. To investigate the molecular determinants of differential replication and pathogenicity, we focused on the polymerase complex as a major contributor to viral replication and pathogenicity ^22,27^. HPhTX and HPbTX polymerases contain three amino acid differences in PB2 (362, 627, and 631), one in PB1 (392), and three in PA (142, 219, and 497) (**Fig. 1)** ^22^. A minigenome (MG) plasmid encoding a double reporter ZsGreen (ZsG) fused to nanoluciferase (Nluc) was generated (**Fig. 2A**). The advantage of this MG encoding ZsG-Nluc is that it allows us to assess polymerase activity by fluorescent microscopy (ZsG) and luminescence (Nluc). Human HEK293T cells were co-transfected with the four plasmids encoding the PB2, PB1, and PA polymerase subunits of HPhTX or HPbTX, together with NP (same amino acid sequence between HPhTX and HPbTX), the MG plasmid, and a pCAGGS plasmid encoding Cypridina luciferase (Cluc) to normalize transfection efficiencies. As internal controls, HEK293T cells were transfected without the PB1 polymerase subunit plasmid. We found that HPhTX has significantly enhanced polymerase activity compared with HPbTX (p= < 0.001) as determined by Nluc activity (**Fig. 2B)** and ZsG expression **(Fig. 2C)**. Notably, all polymerase subunits (PB2, PB1, and PA), and NP were expressed to comparable levels as determined by Western blot **(Fig. 2D)**. These findings indicate that the HPhTX polymerase replicates more efficiently the MG than the HPbTX in HEK293T cells without different levels of polymerase expression.

**Figure 1:**
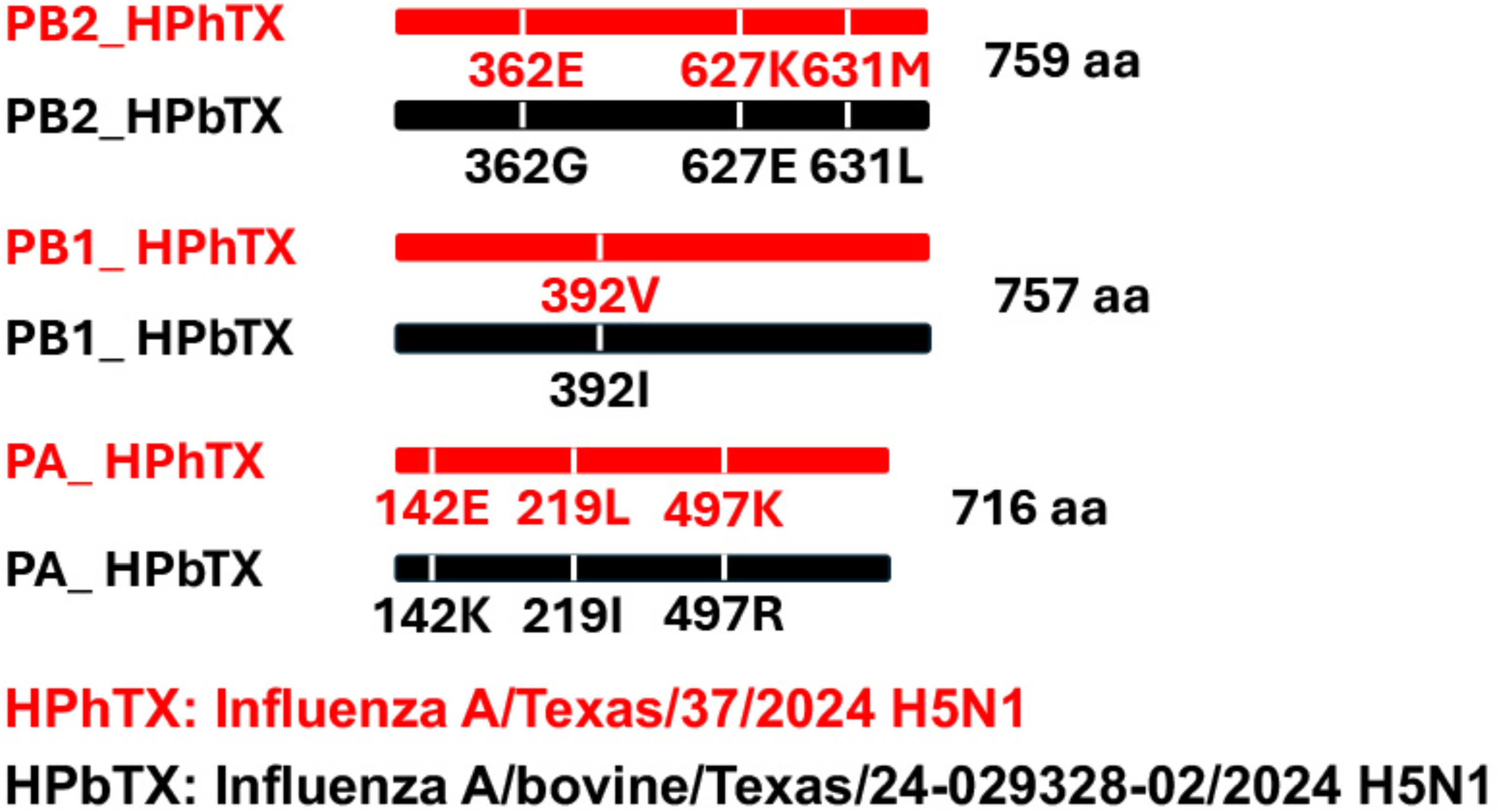
Schematic representation of the amino acid (aa) variations between influenza A/Texas/37/2024 H5N1 (HPhTX, red) and influenza A/bovine/Texas/24-029328-02/2024 H5N1 (HPbTX, black) PB2, PB1, and PA proteins (top to bottom). Protein sizes are indicated in a number on the right.

**Figure 2:**
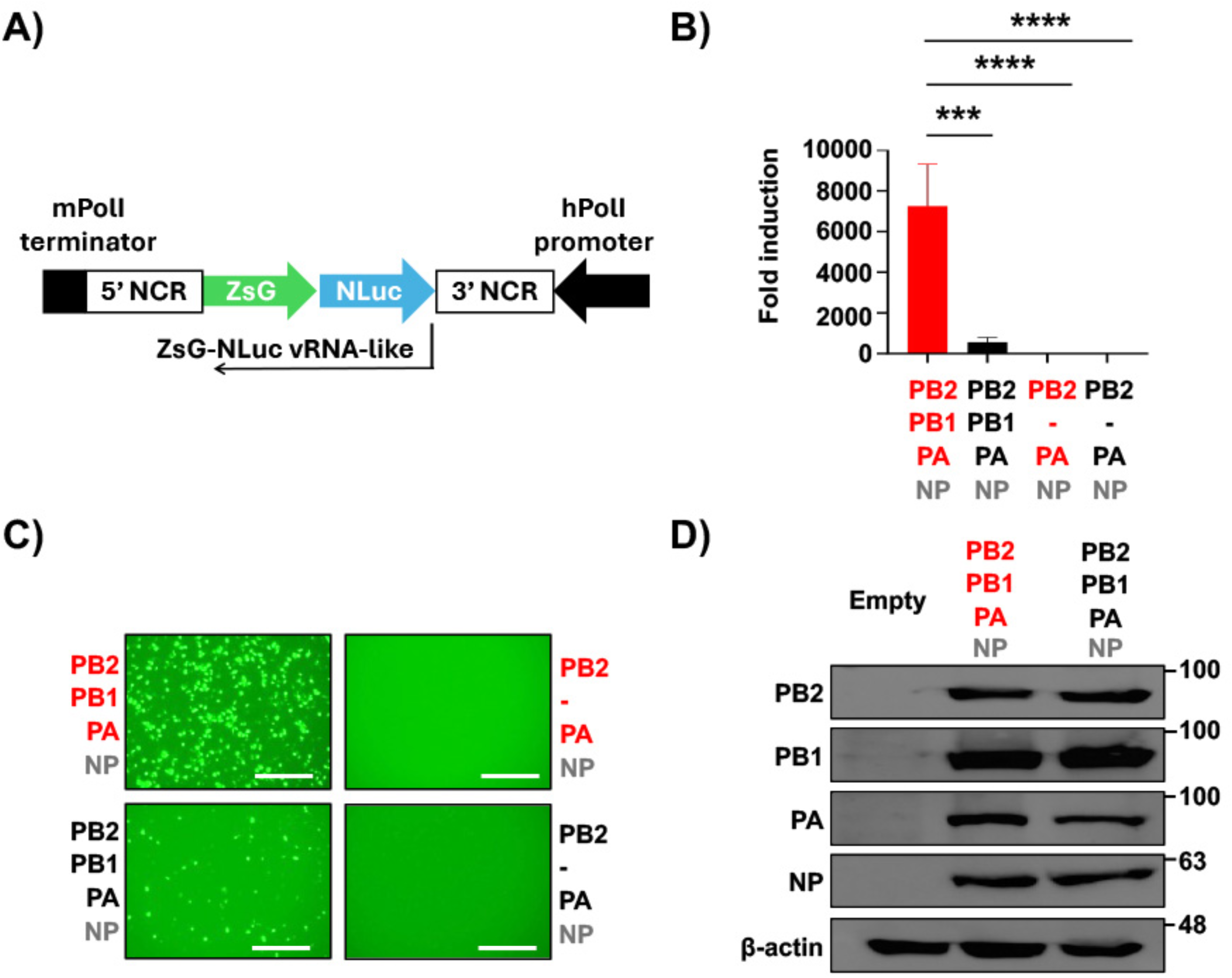
IAV HPhTX has enhanced polymerase activity compared to HPbTX. **A)** Schematic representation of the MG plasmid to assess viral polymerase genome replication and gene transcription. The human polymerase I (hPolI) promoter is indicated as a black arrow. The mouse polymerase I (mPolI) terminator is indicated as a black box. Non-coding regions (NCR) of the HPhTX NP segment are indicated with white rectangles. The ZsG fused to Nluc is indicated. **B)** Cell culture supernatants from HEK293T cells transfected with the MG plasmid, together with the pHW2000 plasmids encoding the polymerase subunits PB2, PB1, PA, and NP of HPhTX (red) and HPbTX (black), and pCAGGS Cluc to normalize transfection efficiency were collected at 30 hpt to assess viral genome replication and gene transcription. Nluc activity was calculated and standardized to Cluc luciferase. HPhTX and HPbTX polymerase activities were expressed as fold induction compared to the negative control lacking the respective pHW2000 PB1 plasmid (-). Data represent the average of three biological replicates with SD indicated. ***p = 0.0001, and ****p < 0.0001 using one-way ANOVA followed by Dunnett’s multiple comparisons test. **C)** Representative images of ZsG expression from cells transfected in (**B**) using live fluorescence microscopy. Scale bars=300 µM. **D)** Western blots: Cell lysates from HEK293T cells transfected in (**B**) were used to assess HPhTX and HPbTX PB2, PB1, PA, and NP expression. Empty pCAGGS plasmid transfected cells served as the negative control. β-actin was included as a loading control. Molecular markers are indicated on the right.

### PB2 segment is responsible for the enhanced polymerase activity of HPhTX

Next, we sought to determine which polymerase subunit(s) was responsible for the increased polymerase activity observed in HPhTX compared to HPbTX. We substituted each of the viral polymerase subunits of the HPhTX with one polymerase segment (PB2, PB1, or PA) of HPbTX (**Figs. 3A and 3B).** Likewise, we substituted each of the viral polymerase subunits of the HPbTX with the PB2, PB1, or PA of HPhTX **(Figs. 3C and 3D)**. The bovine polymerase PB1 and PA subunits exhibited reduced but no statistically significant effect on the human viral polymerase activity in the MG assay **(Figs. 3A and 3B)**. Substituting the human PB2 with the bovine PB2 substantially influenced polymerase activity negatively **(Figs. 3A and 3B)**. Similarly, substituting the human polymerase PB1 and PA subunits did not significantly impact the bovine polymerase activity in the MG assay **(Figs. 3C and 3D)**. However, replacing the PB2 subunit of HPbTX with that of HPhTX significantly enhanced polymerase activity to levels comparable to the parental HPhTX polymerase **(Figs. 3C and 3D)**. These findings demonstrate that the PB2 subunit is responsible for the differences in polymerase activity between the HPhTX and HPbTX in human HEK293T cells.

**Figure 3:**
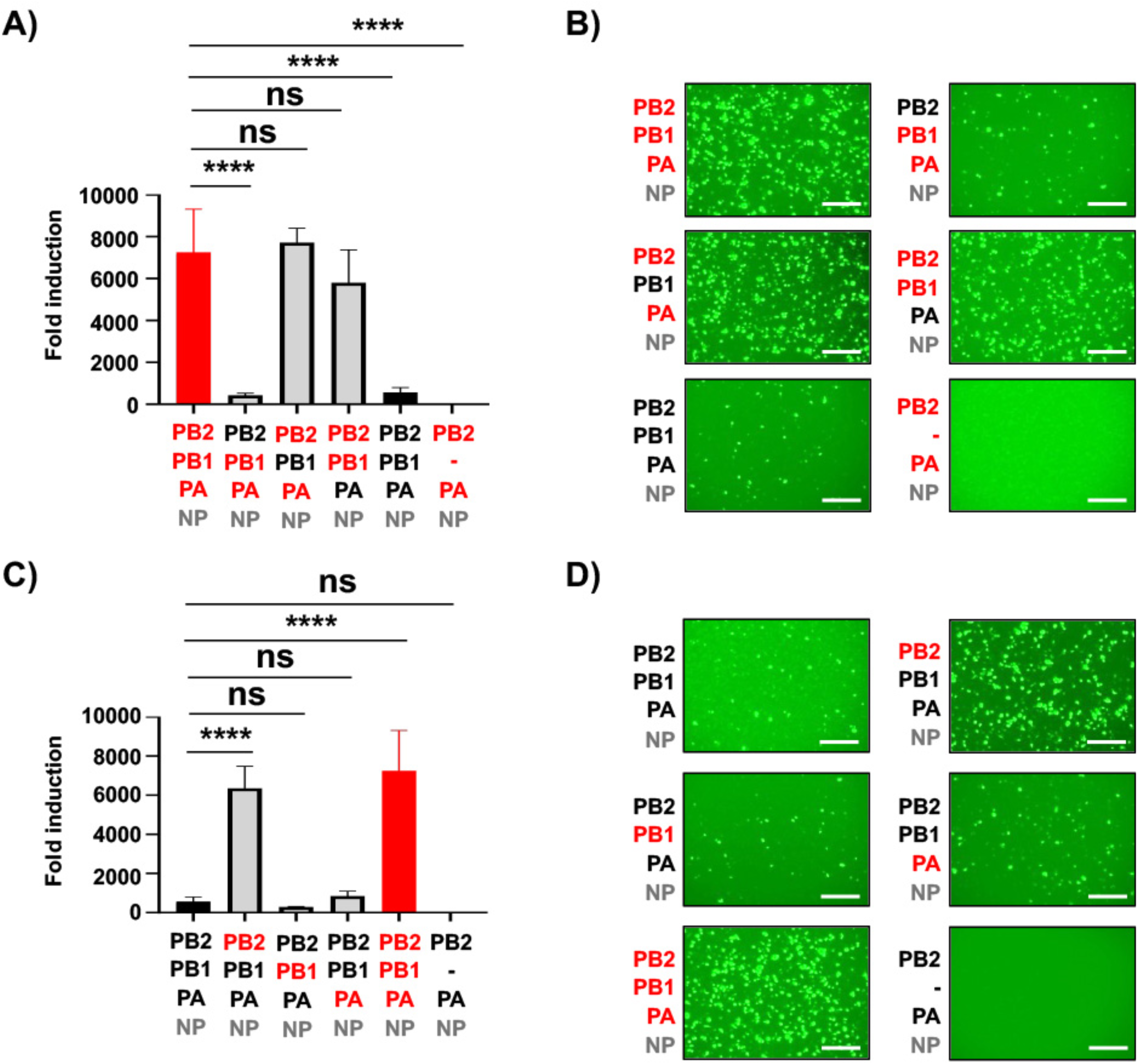
PB2 is responsible for the enhanced polymerase activity of the HPhTX-H5N1. **A and C)** Nluc measurements of the MG assays to evaluate the polymerase activity of HPhTX (**A**) or HPbTX (**C**) 30 h after transfection. Each of the pHW2000 plasmids encoding polymerase subunits of the HPhTX was individually substituted by the PB2, PB1, and PA of the HPbTX (**A**). Likewise, each of the pHW2000 plasmids encoding the polymerase PB2, PB1, and PA subunits of the HPbTX were substituted by the HPhTX counterparts (**C**). Nluc activity was calculated and standardized to Cluc luciferase and expressed as fold induction compared to the negative control sample lacking PB1 (-). Data represent the average of three biological replicates with SD indicated. ns= non-significant and ****p < 0.0001 using one-way ANOVA followed by Dunnett’s multiple comparisons test. **B and D)** Representative images of ZsG expression from cells transfected in (**A**) and (**C**) using live fluorescent microscopy. Scale bars=300 µM.

### Amino acids 362 and 627 in PB2 are responsible for the enhanced polymerase activity of HPhTX

Next, we investigated the amino acid(s) responsible for the enhanced polymerase activity of HPhTX PB2. Three amino acid differences at positions 362, 627, and 631 (**Fig 1**) were identified between HPhTX and HPbTX. We generated HPhTX PB2 plasmids each containing a single amino acid substitution (362G, 627E, and 631L), which is present in the PB2 of HPbTX **(Figs. 4A and 4B)**. Likewise, we generated HPbTX PB2 plasmids containing the individual amino acid substitutions 362E, 627K, and 631M present in the PB2 of HPhTX **(Figs. 4C and 4D)**. We co-transfected HEK293T cells with each of the HPhTX polymerase subunits and the PB2-WT, -362G, -627E, and -631L to assess viral replication and transcription using the MG assay. PB2 362G mutation in HPhTX significantly reduced MG activity **(Figs. 4A and 4B)**. Interestingly, PB2 627E mutation completely abolished polymerase activity **(Figs. 4A and 4B)**. In contrast, PB2 631L demonstrated significantly enhanced (two times) polymerase activity compared to HPhTX PB2 **(Figs. 4A and 4B)**. In the case of the HPbTX MG assays, PB2 HPbTX harboring the 627K mutation restored polymerase activity to levels comparable to HPhTX **(Figs. 4C and 4D)**. PB2 HPbTX 362E mutant also resulted in enhanced polymerase activity, but not to the levels observed with the 627K mutation, while 631M did not significantly change the polymerase activity of PB2 HPbTX WT (**Figs. 4C and 4D**). Western blots confirmed similar protein expression levels for WT and mutant PB2 HPhTX and HPbTX **(Figs. 4A and 4C, respectively)**. These results suggest that amino acid 627 in PB2 is the primary determinant of enhanced polymerase activity of HPhTX, with amino acid 362 also playing a minor, yet significant, role.

**Figure 4:**
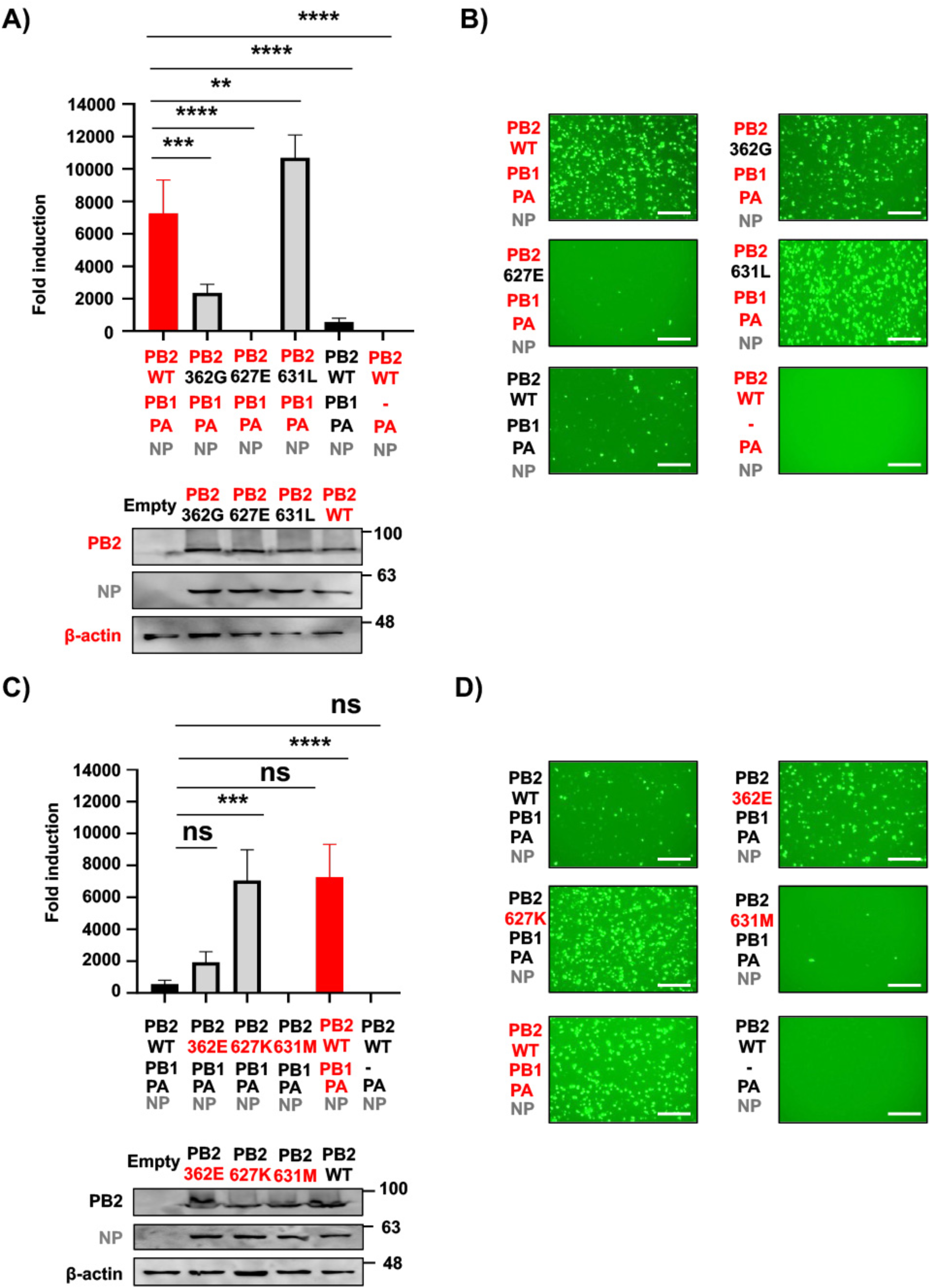
PB2 amino acids 362 and 627 are responsible for differences in polymerase activity between HPhTX and HPbTX. **A and C)** Nluc measurements of the MG assays to evaluate the polymerase activity of HPhTX (**A**) or HPbTX (**C**) 30 h after transfection. The PB2 polymerase subunit of the HPhTX was substituted to a HPhTX PB2 containing mutations 362G, 627E, and 631L (**A**). Likewise, each of the pHW2000 plasmids encoding the polymerase PB2 WT of HPbTX was substituted by HPbTX PB2 containing mutations 362E, 627K, and 631M (**C**). Nluc activity was calculated and standardized to Cluc luciferase and expressed as fold induction compared to the negative control sample lacking PB1 (-). These data represent the average of three biological replicates with SD indicated. ns= non-significant, **p = 0.0067, ***p = 0.0004 and ****p < 0.0001 using one-way ANOVA followed by Dunnett’s multiple comparisons test. Western blots: Cell lysates from HEK293T cells transfected in (**A**) and (**C**) were used to assess PB2 WT and mutant expression levels. Levels of NP expression were included as control. Empty pCAGGS plasmid transfected cells served as the negative control. β-actin was included as a loading control. Molecular markers are indicated on the right. **B and D)** Representative images of ZsG expression from cells transfected in (**A**) and (**C**) using live fluorescent microscopy. Scale bars=300 µM.

Based on these initial results with individual PB2 mutants, we next investigated whether any of these mutations in combination might enhance or reduce viral replication and transcription. To this end, additional PB2 mutants, each harboring two mutations of interest, were generated in the backbone of the HPhTX **(Figs. 5A and 5B)** or HPbTX **(Figs. 5C and 5D)** PB2. Combining the 362G and 627E mutations in the HPhTX PB2 completely abolished polymerase activity, resulting in lower activity than the WT HPbTX polymerase **(Figs. 5A and 5B)**. Combining the 362G and 631L mutations in the HPhTX PB2 did not significantly affect the viral polymerase activity **(Figs. 5A and 5B)**. Conversely, the presence of 627E and 631L mutations in the human PB2 significantly reduced polymerase activity compared to the HPbTX PB2 **(Figs. 5A and 5B)**. These results further confirm our previous results with individual mutants suggesting that amino acid 627 is the primary determinant of the differences in viral replication and transcription between the HPhTX and the HPbTX **(Fig. 4)**. Likewise, PB2 362E and 627K mutations in the HPbTX PB2 resulted in significantly enhanced polymerase activity, exceeding that of WT HPhTX **(Figs. 5C and 5D)**. PB2 362E and 631M did not significantly affect the polymerase activity of HPbTX **(Figs. 5C and 5D)**. Double 627K and 631M mutation in HPbTX PB2 restored, to some extent, the polymerase activity but not to levels observed in HPhTX PB2 **(Figs. 5C and 5D)**. Western blot confirmed similar WT and mutant PB2 protein expression levels of the HPhTX and the HPbTX (**Figs. 5A and 5C, respectively**). These findings indicate that PB2 E627K mutation is responsible for the major differences in polymerase activity between HPhTX and HPbTX and that G362E mutation, together with E627K, plays a synergistic role in enhancing the polymerase activity of HPbTX in human cells. Likewise, the M631L mutation in HPbTX PB2 contributes to enhanced polymerase activity in conjunction with E627K in HEK293T cells.

**Figure 5:**
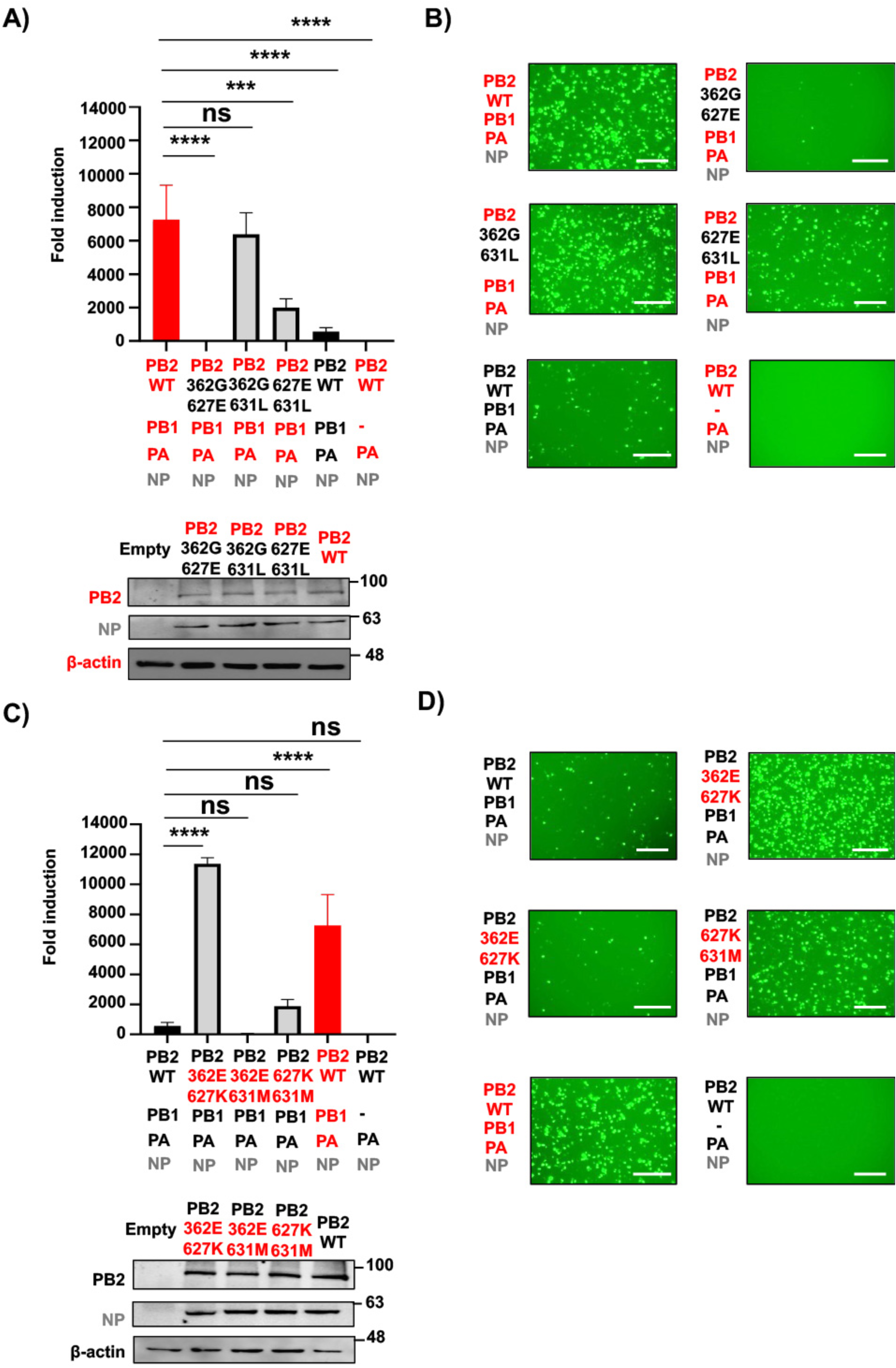
Amino acid substitution 362G, 627E, and 631M in PB2 are responsible for the reduced polymerase activity of HPbTX. **A and C)** Nluc measurements of the MG assays to evaluate the polymerase activity of HPhTX (**A**) or HPbTX (**C**) 30 h after transfection. The PB2 polymerase subunit of the HPhTX was substituted to PB2 containing the combination of mutations 362G/627E, 362G/631L, and 627E/631L (**A**). Likewise, each of the pHW2000 plasmids encoding the polymerase PB2 WT of HPbTX was substituted by HPbTX PB2 containing the combination of mutations 362E/627K, 362E/631M, and 627K/631M (**C**). Nluc activity was calculated and standardized to Cluc and expressed as fold induction compared to the negative control sample lacking PB1 (-). These data represent the average of three biological replicates with SD indicated. ns= non-significant, **p = 0.0067, ***p = 0.0004 and ****p < 0.0001 using one-way ANOVA followed by Dunnett’s multiple comparisons test. Western blots: Cell lysates from HEK293T cells transfected in A and C were used to assess HPhTX and HPbTX polymerase PB2 WT and mutant expression levels. Levels of NP expression were included as control. Empty pCAGGS plasmid transfected cells served as the negative control. β-actin was included as a loading control. Molecular markers are indicated on the right. **B and D)** Representative images of ZsG expression from cells transfected in (**A**) and (**C**) using live fluorescent microscopy. Scale bars=300 µM.

### Generation and characterization of rHPhTX-PB2 K627E, -PB2 E362G, and -PB2 E362G/K627E viruses

The above data indicate that PB2 mutations K627E and E362G are the major factors responsible for the differences in polymerase activity between the HPhTX and HPbTX. Building upon these previous results, we generated, using our previously described HPhTX reverse genetics approaches ^22^, rHPhTX containing K627E, E362G, or both K627E/E263G mutations in PB2 to assess the contribution of these amino acid changes in viral pathogenicity using a loss of function (LoF) approach. NGS sequencing of the full-length PB2 segment confirmed the successful rescue of these mutant rHPhTX **(Fig. 6A)**. Next, we performed replication kinetic analyses in A549, a human cell line distinct from the HEK293T cells used in the MG assays, bovine MDBK, and canine MDCK cells. Results consistently demonstrate significant enhanced viral replication of rHPhTX, particularly evident after 24 hpi, aligning with the findings in our MG assays **(Figs. 6B-6D)**. Interestingly, enhanced replication was found not only in human cells, but also in bovine and canine cells. The plaque phenotype of the WT and mutant rHPhTX in the same cell lines confirmed the competent replication of all viruses **(Figs. 6B-D**) and suggested that both PB2 mutations reduce rHPhTX replication.

**Figure 6:**
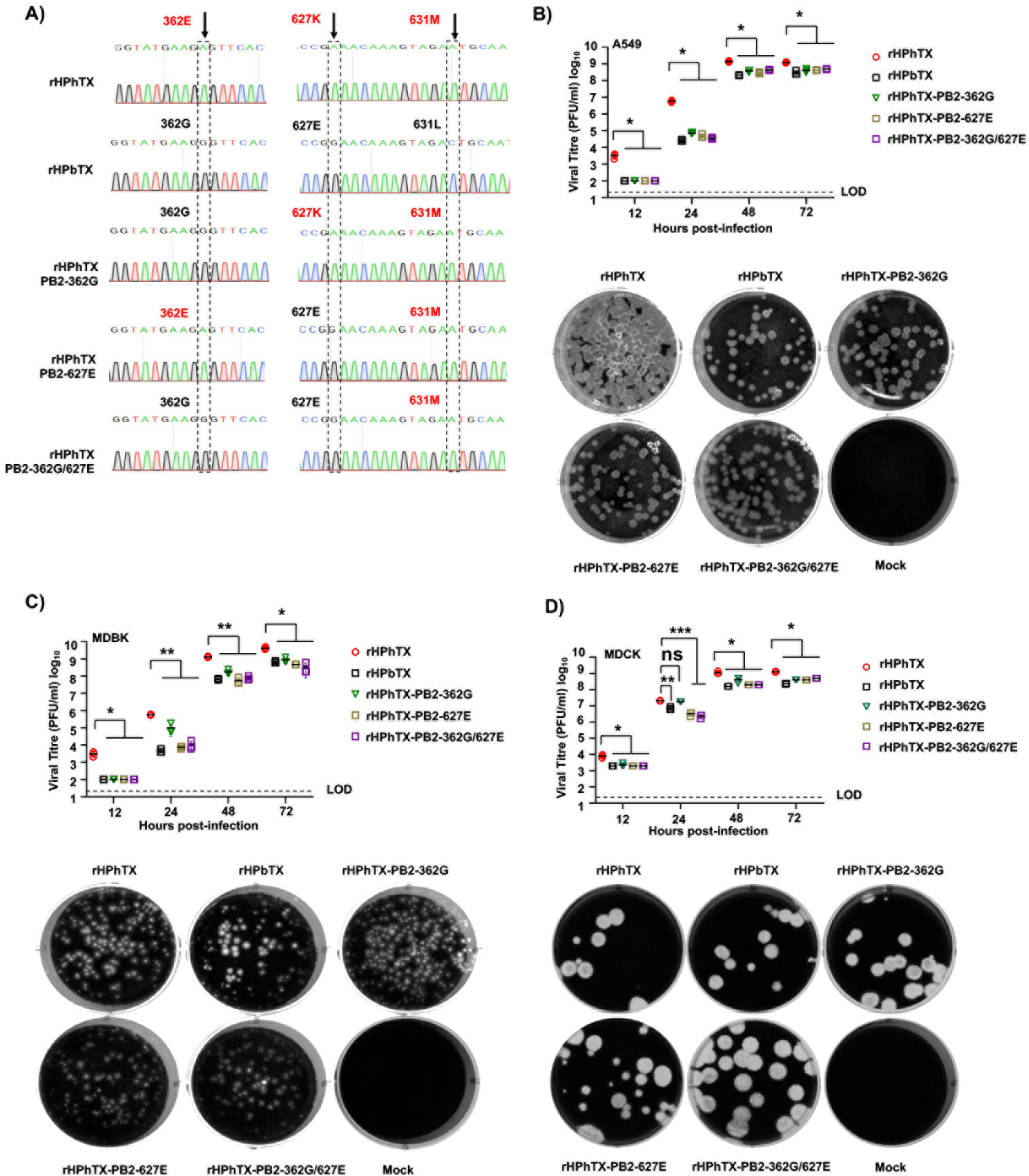
Generation and *in vitro* characterization of rHPhTX PB2-K627E, -E362G, and -E362G/K627E. **A)** NGS of the PB2 segment from rHPhTX PB2-K627E, -E362G, and -E362G/K627E generated using reverse genetics to confirm the presence of mutations. The location of amino acid residues 362, 627, and 631 in the different recombinant viruses is shown. **B-D)** Viral growth kinetics and plaque assay of WT and mutant PB2 rHPhTX and rHPbTX in A549 **(B)**, MDBK **(C)**, and MDCK **(C)** cells. Virus titration was performed using standard plaque assays on MDCK cells, followed by crystal violet staining. Data represent the average of three biological replicates with SD indicated. ns= non-significant, *p < 0.01, **p < 0.001 and ***p < 0.0001 using two-way ANOVA with Greenhouse-Geisser correction, followed by Tukey’s multiple comparisons analysis. The limit of detection (LoD, 20 PFU) is indicated.

To assess the impact of these mutations on viral pathogenicity, we infected 6-week-old female C57BL/6J mice with 20 plaque-forming units (PFU) of the WT and mutant PB2 rHPhTX. As anticipated, rHPhTX WT caused rapid weight loss and 100% mortality by day 8 **(Figs. 7A and 7B)**. Interestingly, rHPhTX PB2 362G also resulted in 100% mortality, albeit a delay in body weight loss and mortality was observed compared to rHPhTX WT **(Figs. 7A and 7B)**. In contrast, rHPhTX PB2 627E and rHPhTX PB2 362G/627E infected mice exhibited only 20% mortality, similar to rHPbTX, suggesting reduced pathogenicity of rHPhTX containing the K627E mutation **(Figs. 7A a d 7B)**. To correlate body weight and survival with viral titers in infected mice at 2 and 4 DPI, nasal turbinate, lungs, and brains from infected mice were collected, homogenized, and subjected to plaque assay on MDCK cells. As expected, rHPhTX WT exhibited the highest viral load in all tissues and days, followed by rHPhTX-PB2-362G **(Fig. 7C)**. Notably, rHPhTX PB2 627E and rHPhTX PB2 362G/627E infected mice showed comparable low viral loads to those infected with rHPbTX **(Fig. 7C)**. These findings confirmed the attenuation achieved through the identified mutations 362G, 627E, and both 362G/627E in PB2.

**Figure 7:**
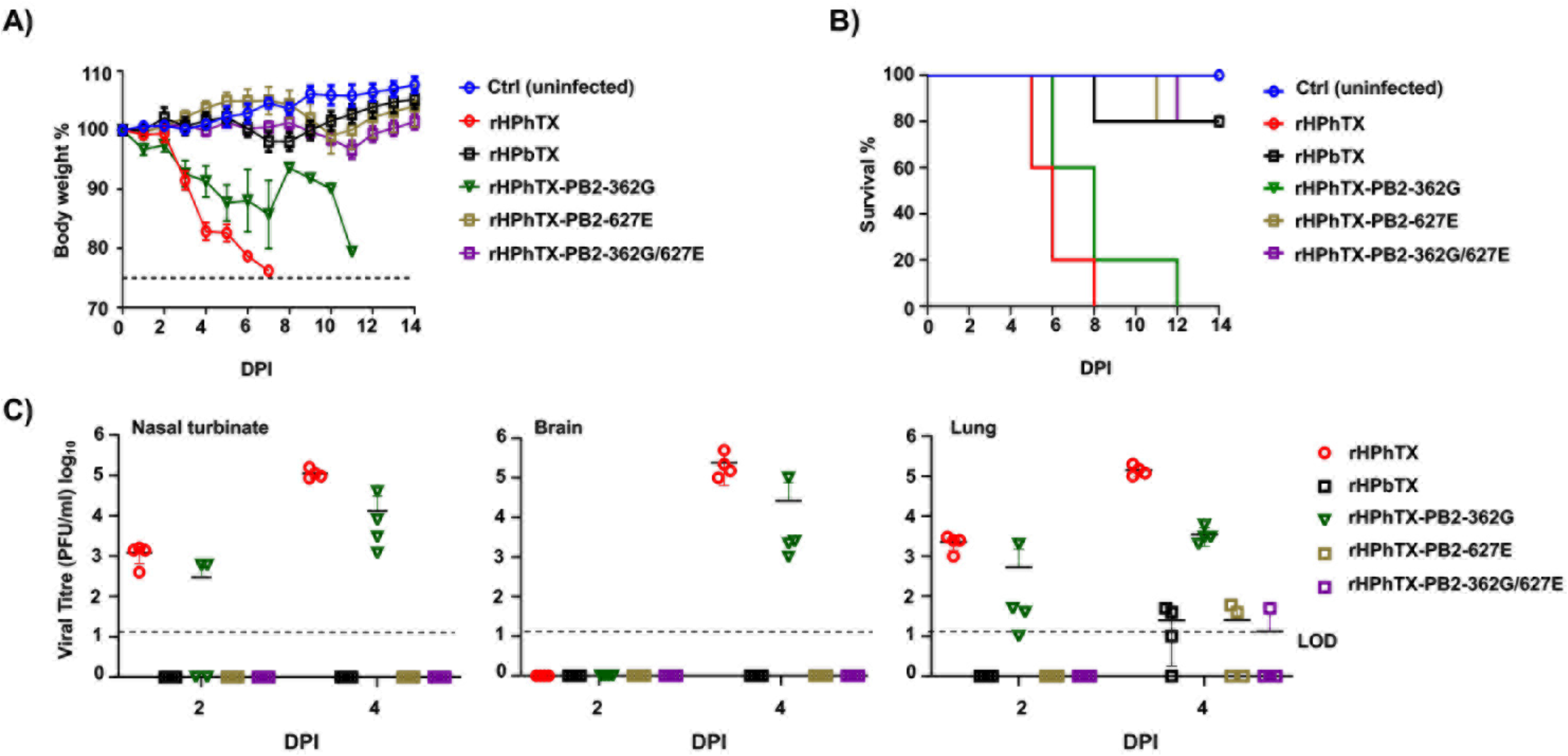
Pathogenicity of rHPhTX PB2-K627E, -E362G, and-E362G/K627E mutants in C57BL/6J mice. **A and B)** Percentage (%) of body weight changes (**A**) and survival curves (**B**) of C57BL/6J mice (n=5/group) infected with 20 PFU of each of the indicated WT or mutant viruses. The mean percent of body weight change (± SD) is indicated. Mice were humanely euthanized when they had lost more than 25% of their initial body weight. **C)** Viral titers in different tissues of 6-week-old female C57BL/6J mice infected with rHPhTX, rHPbTX, rHPhTX PB2-K627E, rHPhTX PB2-E362G, and rHPhTX PB2-E362G/K627E viruses. Nasal turbinate, brains and lungs were collected at 2 and 4 DPI (n=4/timepoint), homogenized, and titrated using standard plaque assays on MDCK cells using crystal violet staining. The limit of detection (LoD, 20 PFU) is indicated.

### Differential pathology of WT and rHPhTX PB2 mutants in murine lungs and brains

In addition to viral replication, we also examined lung and brain histopathology in C57BL/6J mice infected with rHPhTX WT and PB2 mutant viruses at 4 DPI. Histopathology analysis and virus-induced pathology were quantified following hematoxylin and eosin (H&E) staining. Serial sections were used for immunohistochemistry (IHC) staining for viral NP antigen using an anti-NP antibody raised in rabbits. Histopathological examination revealed that rHPhTX WT infection caused multifocal to extensive alveolar septal necrosis, with moderate-to-severe infiltration of lymphocytes and plasma cell infiltrates, along with edema, hemorrhage, fibrin, and cellular debris (average percent of lung pathology of 40.4%) **(Figs. 8A and 9)**. Affected bronchioles showed epithelial necrosis and replacement by cellular and karyorrhectic debris **(Fig. 8A)**. In contrast, rHPbTX-infected mice exhibited minimal lymphocytic infiltration around blood vessels and bronchioles, both consistent with our previous report ^22^ **(Fig. 8A)**. Infection with rHPhTX PB2 362G resulted in milder (∼4X lower) lesions than rHPhTX WT-infected mice (average percent lung pathology of 10.8%), characterized by multifocal lesions around bronchioles and blood vessels, extending to alveolar septa, with moderate lymphocyte, plasma cell, and macrophage infiltrates **(Figs. 8A and 9)**. Affected bronchi and bronchioles showed attenuation, ciliary loss, epithelial necrosis, and replacement by cellular and karyorrhectic debris mixed with inflammatory cells. Mild edema and hypertrophied endothelium with inflammatory cell accumulation within the tunica intima were also observed **(Fig. 8A)**. Infection with rHPhTX PB2 627E and rHPhTX PB2 362G/627E caused very mild lesions in only one mouse per group (average percent lung pathology less than 1%), including segmental bronchial epithelial loss and small infiltrates of inflammatory cells and necrotic debris, similar to mice infected with rHPbTX (average percent lung pathology less than 1%). Brain tissues showed negligible histopathological changes across all groups **(Fig. 8B)**.

**Figure 8:**
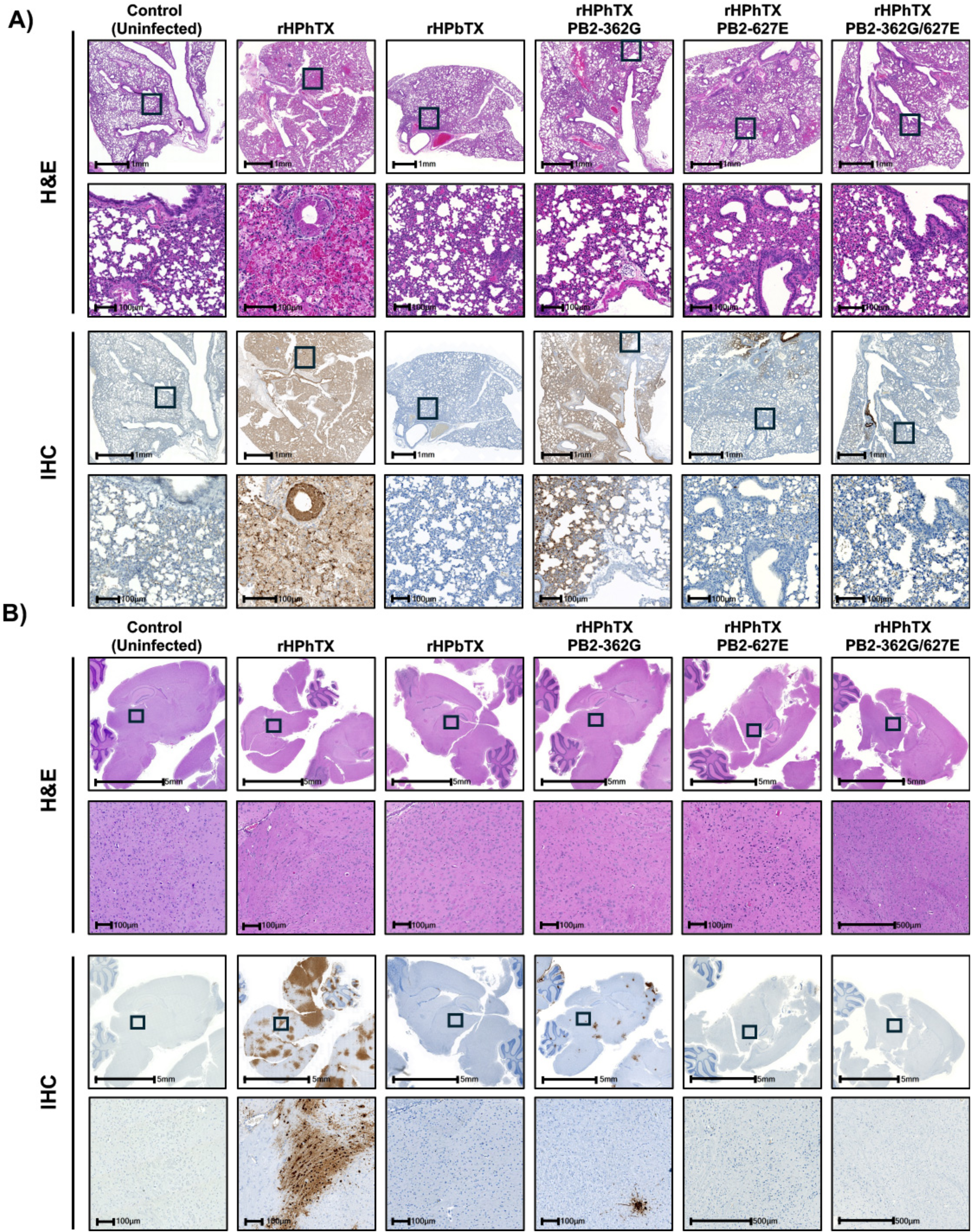
Histopathological alterations in the lungs and brains of C57BL/6J infected mice. Histopathological alterations in the lung **(A)** and brain **(B)** tissues of mice infected in Figure 7 compared to uninfected control C57BL/6J mice. The upper 2 rows display the tissue stained with hematoxylin-eosin (H&E). The zoom-in area in the lower images is indicated by black boxes on the top images. The bottom 2 rows show the IHC results for the same tissue sections, stained with a rabbit polyclonal antibody against the viral NP. Scale bars: the upper row in both H&E and IHC scale bars = 1 mm, whereas the zoom-in area = 0.1mm **(A)**, the upper row in both H&E and IHC scale bars = 5 mm, whereas the zoom-in area = 0.1mm **(B)**.

**Figure 9:**
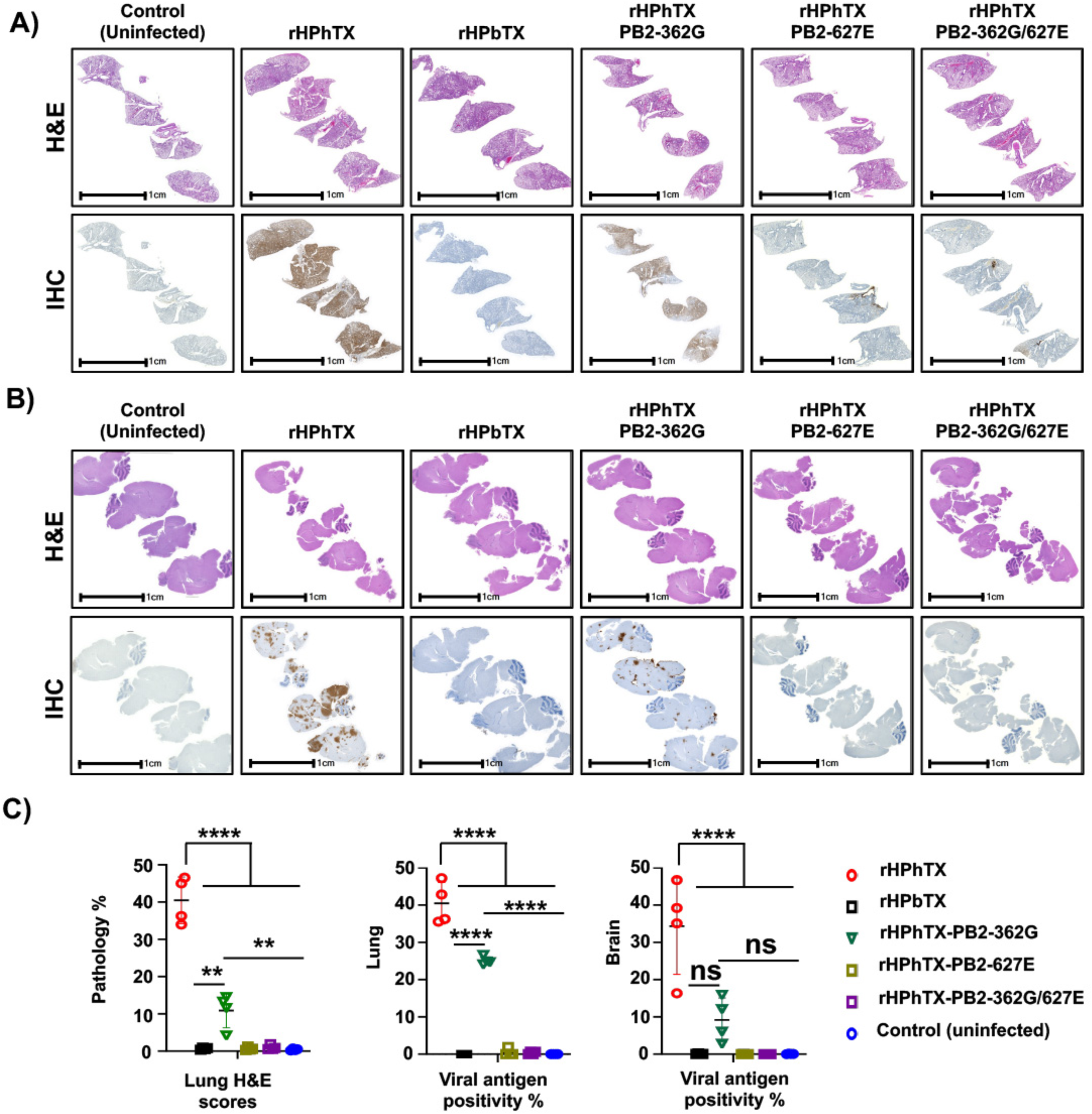
Histopathological alterations, lesion scores, and % of viral NP antigen in the lungs and brains of infected C57BL/6J mice. **A and B)** Histopathological alterations in the lung (**A**) and brain (**B**) tissues of mice infected with the indicated viruses compared to uninfected mice from tissues collected in Figure 7. The upper images displayed in each panel show tissue stained with hematoxylin-eosin (H&E) for each virus. The bottom images show the immunohistochemistry (IHC) results for the same tissue sections, stained with a rabbit polyclonal antibody against the viral NP. **C)** A quantitative evaluation of H&E lesion scores and tissue viral NP antigen (%) in the lung and brain tissues of mice infected with the indicated viruses compared to the uninfected group. The overall pathology score was established based on virus quantification in the lung and brain tissues of the infected C57BL/6J mice (n=4) from Figure 7. The data are expressed as mean ± SD. Statistical comparisons among the indicated groups were conducted using one-way ANOVA followed by Dunnett’s multiple comparisons. Significant differences are denoted as follows: ** = p < 0.01, *** = p < 0.001, **** = p < 0.0001; non-significant = ns. Scale bars are indicated, and each equal 1 cm.

Immunohistochemical staining revealed positive NP viral antigen staining in lungs and brains, varying significantly between viruses, with the most extensive staining in rHPhTX-infected mice (the average percentage of lung and brain staining for viral antigen was ∼40% and 34%, respectively), followed by rHPhTX PB2-362G (i.e., 25% and 9%, respectively; **Figs. 9A and B)**. Mice infected with rHPbTX, rHPhTX PB2-627E, and rHPhTX PB2-362G/627E groups showed negligible viral antigen (average percentage of lung staining for viral antigens is less than 1%) **(Figs. 9A and B)**. Lungs and brains from control mock-infected mice exhibited normal epithelium, no significant histopathological changes, and negative IHC **(Figs. 8 and 9)**.

## Discussion

IAVs pose serious public health problems, causing severe seasonal epidemics and occasional pandemics of great concern to humans. Two critical factors in the evolution of IAVs significantly contribute to their widespread prevalence. First, novel IAVs can emerge unexpectedly in avian reservoirs through genetic reassortment or direct mutations, potentially acquiring the ability to infect and transmit in a naïve human population and initiating a pandemic. Second, after infection in humans, IAVs can undergo rapid selection and acquire unpredictable changes in their antigens ^1,2^. A recent notable example of this phenomenon is the H5N1 outbreak in the US, which involved spillover events from wild birds to dairy cattle and, subsequently, to humans ^15,27,30^. The first human H5N1 isolate from a dairy cattle worker was reported in Texas in April 2024 ^20^. Our earlier investigation revealed enhanced replication kinetics and pathogenicity of this human strain (HPhTX) compared to the closely related cattle strain (HPhTX) ^22^. Herein, we reveal that HPhTX exhibits enhanced polymerase activity compared to HPbTX in HEK293T cells, primarily due to the PB2 polymerase subunit. We also demonstrate that mutations G362E and E627K are the major contributors to the differences in polymerase activity between the HPhTX and HPbTX strains. Using reverse genetics and a LoF approach, we also demonstrate the contribution of these amino acids at positions 362 and 627 as responsible for enhanced pathogenicity of rHPhTX as compared to rHPbTX in C57BL/6J mice.

It has been previously shown that polymerase activity is an important pathogenicity factor of IAVs ^22,27^. Thus, we sought to determine if the enhancement in viral replication and pathogenicity in mice stemmed primarily from differences in the polymerase complex between the human and bovine isolates. Notably, 7 out of the 9 amino acid differences between HPhTX and HPbTX isolates located within the polymerase PB2, PB1, and PA subunits **(Fig. 1)**. Using a novel MG plasmid expressing a fusion of ZsG and Nluc, we demonstrated a marked increase in polymerase activity of the human polymerase complex in human cells, corroborating the previously observed enhanced replication of HPhTX **(Fig. 2)** ^22^.

Swapping individual subunits between human and bovine polymerase subunits revealed PB2 as the primary determinant of the differences in polymerase activity **(Fig. 3)**. Mutations in other polymerase subunits (i.e., PB1 and PA) did not show a significant effect on viral replication in the MG assay **(Fig. 3)**. Since only three distinct amino acid differences were present between the human and bovine PB2, we next analysed which one(s) were responsible for the differences in viral replication in HEK293T cells. Analysis of single and double PB2 substitution revealed that PB2 E627K is the major responsible for the observed differences in polymerase activity **(Figs. 4 and 5)**. Notably, this mutation is a key marker frequently associated with avian-to-mammalian host adaptation, facilitating productive viral replication and infection ^31–33^. More interestingly, the M631L mutation in HPhTX PB2 was responsible for an enhancement in polymerase activity **(Figs. 4 and 5)**. M631L substitution has been previously shown to be another marker of mammalian adaptation that supports productive infection after spillover from wild-bird reservoirs ^27,34^. These findings are corroborated by a recent study on the same isolate identified in humans ^27,34^. However, this previous report did not address the impact of other mutations on the polymerase complex in human polymerase activity, viral replication, and pathogenicity. Our MG results also show that G362E plays an important role in human polymerase activity **(Figs. 4 and 5)**. To investigate the effects of these mutations on pathogenicity, we rescued mutant viruses using a loss-of-function (LoF) approach **(Fig. 6)**. We generate rHPhTX containing mutations in PB2 affecting polymerase activity (i.e., E362G, K627E, and E362G/K627E) **(Fig. 6)**. While all these viruses replicated in human (A549) cells, they showed reduced replication compared to rHPhTX WT **(Fig. 6)**. Interestingly, the same phenotype was obtained in bovine (MDBK), and canine (MDCK) cells. Importantly, none of the rHPhTX viruses containing mutations in 362 and 627, alone or in combination, replicated more efficiently than the rHPhTX in any of the investigated cell lines, demonstrating the LoF associated with these mutations **(Fig. 6)**.

To investigate the role of PB2 mutation in viral pathogenicity, we infected 6-week-old female mice with 20 PFU **(Fig. 7)**. We previously showed that mice infected with 10 PFU of the rHPhTX rapidly lost weight, and all succumbed to viral infection, contrary to mice infected with 10 PFU of the rHPbTX, where about 80% survived viral infection ^22^. While rHPhTX PB2 362G exhibited the same mortality rate as rHPhTX, infected mice showed a delay in body weight loss and mortality **(Fig. 7)**. This finding indicates that PB2 E362G mutation results in viral attenuation compared to rHPhTX, supporting our findings in the MG assay **(Fig. 4)**. Interestingly, rHPhTX PB2 627E and rHPhTX PB2 362G/627E showed similar body weight and mortality rates in mice than rHPbTX (20%) ^22^. Viral titers and staining of viral antigens using IHC in mice lungs and brains corroborate these findings **(Figs. 8 and 9)**. Notably, rHPhTX PB2 627E and rHPhTX PB2 362G/627E had minimal neurotropic lesions and tissue pathology scores **(Figs. 8 and 9)**. However, mice infected with rHPhTX PB2 362G had a moderate-to-severe impact on mice brains **(Figs. 8 and 9)**. These findings suggest that the 627 mutation in PB2 is primarily responsible for the differences in replication kinetics and pathogenicity of rHPhTX compared to rHPbTX and that the 362 residue has a minor effect.

IAV PB2 subunit consists of different domains with amino acids 362, 627, and 631 located in the cap-binding (362) and 627 domain (627 and 631) **(Fig. 10A)**. Previous studies have shown that phenylalanine residues at positions 363 and 404 are crucial for the effective cap-binding activity of IAV due to their conserved aromatic rings ^35^. Given that the 362 residue is in close proximity to these two aromatic residues, the presence of 362G may slightly increase the distance between these aromatic rings in the predicted structure of bovine PB2 when compared to the human PB2 **(Fig. 10B)** and affect the PB2 cap-binding activity. However, obtaining the crystal structure of the bovine H5N1 PB2 and, more importantly, the polymerase complex, is essential to validate this hypothesis. Additionally, another amino acid 368 mutation in PB2 located near amino acid 362 has been documented to influence the pathogenicity of H5N1 in mice ^36^. Notably, avian-like arginine (R) residue at 368, found in both HPhTX and HPbTX, could affect cap binding ability, warranting further investigations ^36^. Remarkably, the 627 and 631 mutations occur in the flexible auxiliary regions of the PB2 and do not contribute to the core polymerase function (**Fig. 10A)**. Importantly, the 627 domain plays an important role in host range restriction of IAVs ^37^. Avian IAV polymerases exhibit limited replication efficiency in mammalian cells and necessitate specific adaptive mutations to restore their polymerase activity. These mammalian host adaptations often involve mutations within, but are not limited to, the 627 domain in PB2. The spillover event from an avian-origin strain linked to the ongoing H5N1 outbreak, specifically A/Canada_goose/Wyoming/24-003692-001-original/2024 (cgWY001-H5N1), was believed to be the parental avian strain, which possesses the 627E and 631M markers of mammalian host adaptation. Following the spillover, adaptive mutations occurred that enabled H5N1 replication in cattle (which adapt to harbour 627E and 631L in the PB2) and the further spillover event of H5N1 to humans (which adapt to harbour 627K and 631M in the PB2) ^27^. A recent investigation has revealed that 631L engages with bovine ANP32A, an essential host factor for IAV replication in mammals ^38^.

**Figure 10:**
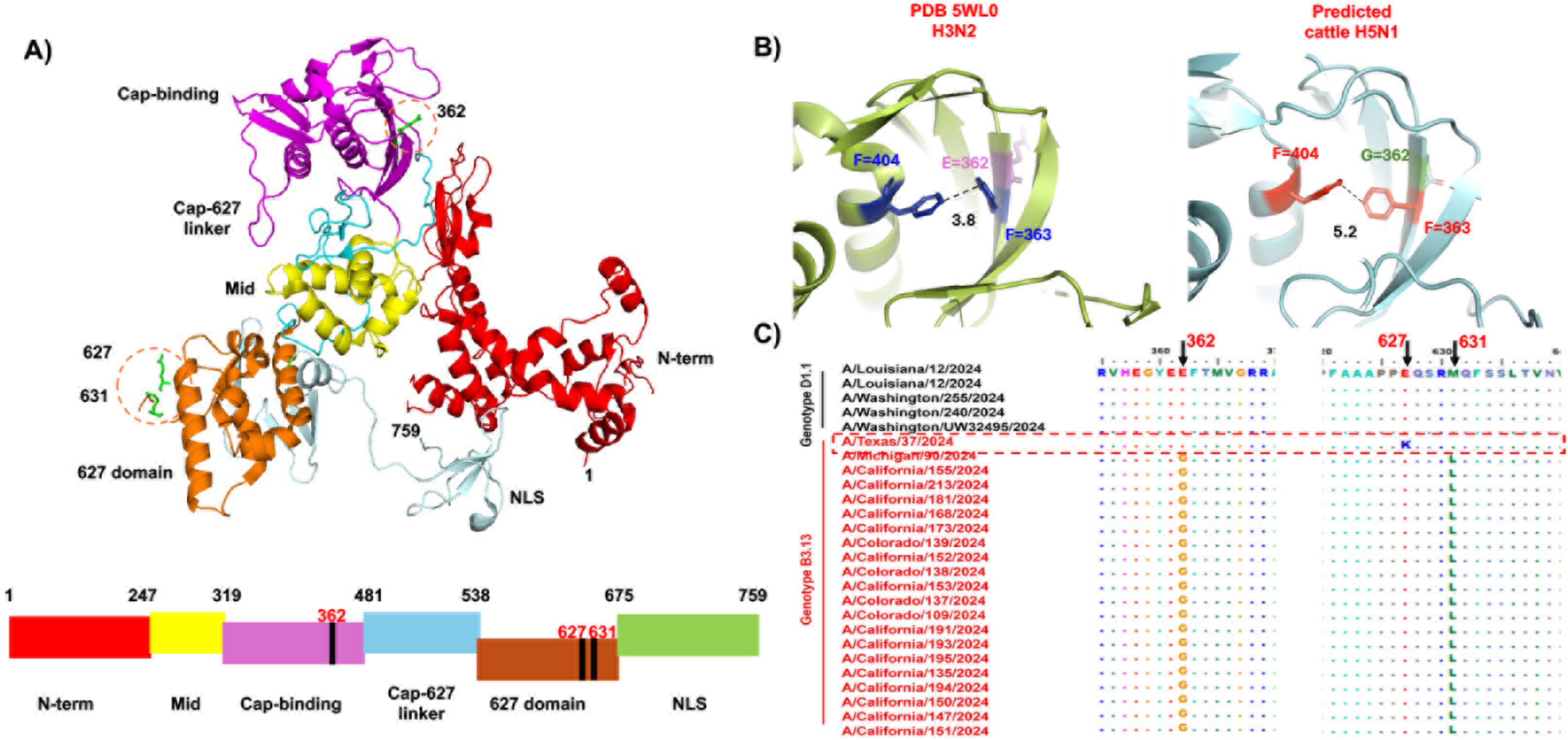
Protein structure of PB2 and the structural differences between HPhTX and HPbTX. **A)** Ribbon 3D diagram of PB2 and its subdomains (PDB: 6QNW A/duck/Fujian/01/2002 (H5N1)). The three different amino acids between HPhTX and HPbTX PB2 are indicated by orange dashed circles. Each subdomain is colored as in the 2D schematic diagram, and shown at the bottom are the 362, 627, and 631 amino acids of interest, which are indicated by black bars. **B)** Structural differences between human (PDB: 5wl0; A/Udorn/307/1972 (H3N2)) and predicted A/bovine/Texas/24-029328-02/2024 H5N1. Cap-binding domains are indicated. The two phenylalanine residues forming the aromatic ring in the cap-binding domain, F363 and F404, and the distance between them are indicated. E362 (human) and G362 (bovine) are indicated by magenta and green, respectively. **C)** Amino acid sequence alignment of genotype D1.1 (black) and B3.13 (red) PB2 proteins of recent H5N1 isolated in the US. The alignment was generated using the ClustalW algorithm of the MegAlign program (DNAstar) and visualized by BioEdit software. The sections containing the mutations of interest are shown. HPhTX is shown by a dashed-red box.

Analysis of HPAIV H5N1 sequences identified in humans during the recent outbreak in the US reveals that the mutation 627K observed in the PB2 gene is distinctive to the HPhTX isolate, not present in other bovine or human B3.13 or D1.1 strains **(Fig. 10C)**. It is also important to highlight that the recently reported human isolates of D1.1 strains, which exhibit enhanced pathogenicity, possess the 362E mutation (and lack 627K) in the PB2 gene. Nevertheless, the molecular factors contributing to their pathogenicity require further investigations ^39^. This data suggests the possibility that these mutations rapidly emerged after human infection with the bovine HPAIV H5N1 for efficient viral replication. Notably, it has been recently found that the HPhTX can also efficiently infect and transmit in ferrets ^27,34^, pointing out that a few amino acid mutations would allow efficient viral replication and transmission among mammals. Furthermore, these results also demonstrate the importance of monitoring HPAIV H5N1 infecting humans for the presence of these or similar mutations that allow them to better replicate and possibly transmit. Finally, these results suggest the importance of active measures in place to eliminate HPAIV H5N1 from cattle to prevent further human infections and potential transmissions of HPAIV H5N1.

## Acknowledgements

We express our gratitude to the Histology unit team members Renee Escalona and Colin Chuba and the entire L.M-S group at Texas Biomed for their valuable assistance. We extend our gratitude to BEI Resources for providing antibodies against H5N1 polymerases. Furthermore, we would like to thank Professor Daniel Perez for providing MDBK cells.

## Authors contributions

Conceptualization: M.B., A.M. and L.M-S.; Methodology: M.B., R.B., R.E., V.S., N.J., C.Y.; Data collection and interpretation: M.B., R.B., R.E., V.S., N.J., C.Y., A.E., L.M-S; Funding acquisition and resources: A.G-S., and L.M-S.; Writing—original draft preparation: M.B., L.M-S; Writing—review and editing: all authors have read and agreed to the published version of the manuscript.

